# Upregulation of CREB1 and FOXO1 transcription factor pathways in Neuregulin-1 mediated neuroprotection following ischemic stroke

**DOI:** 10.1101/2021.11.17.468955

**Authors:** Kimberly R. Bennett, Monique C. Surles-Zeigler, Catherine J. Augello, Etchi Ako, Victor G. J. Rodgers, Byron D. Ford

## Abstract

Neuregulin-1 (NRG-1) is growth factor that has been investigated for its neuroprotective properties following ischemic stroke. While NRG-1 has shown significant promise in preventing neuronal damage following stroke, the mechanisms behind its neuroprotective effects are unclear. The goal of this research was to investigate the effects of NRG-1 treatment on ischemia-induced gene expression profiles following a permanent middle cerebral artery occlusion (MCAO) in rats. Rats were sacrificed twelve hours following MCAO and either vehicle or NRG-1 treatment. RNA extracted from the peri-infarct cortex of the brain was hybridized to an Affymetrix Rat Genome 2.0st Microarray Gene Chip. Data were analyzed using the Affymetrix Transcriptome Analysis Console (TAC) 4.0 software and the STRING Protein-Protein Interaction Networks database. Our results showed that NRG-1 delivery increased the regulation of pro-survival genes. Most notably, NRG-1 treatment upregulated the CREB1 and FOXO1 transcription factor pathways which are involved in increasing anti-inflammatory and cell proliferation responses and decreasing apoptosis and oxidative stress responses, respectively. Luminex multiplex transcription factor assays demonstrated that the activities of CREB1 and FOXO1 were increased by NRG-1 treatment with MCAO. These findings provide novel insight into the molecular mechanisms involved in NRG-1 mediated neuroprotection.

## Introduction

Stroke is a leading cause of death worldwide, with about 87% of cases being ischemic stroke [1-3]. Differing from hemorrhagic stroke, which occurs due to cerebrovascular blood loss, ischemic stroke occurs when the brain blood supply is obstructed. Ischemic stroke is characterized by cell death, inflammation, and oxidative stress, that occur through molecular cascades and changes in regulatory gene expression [4-6]. The only approved treatment for ischemic stroke is tPA, a limited and time-sensitive treatment aimed at restoring blood flow that only 3-5% of patients qualify for [1, 2]. Given the wide-spread and dire need for an effective ischemic stroke treatment that can be administered outside the 3-hour therapeutic window of tPA, alternate therapies are needed.

One potential therapy for stroke is delivery of neuregulin-1 (NRG-1). NRG-1 has been investigated as a potential therapeutic for ischemic stroke for its various roles in the nervous system [7-15]. NRG-1 is a small protein first discovered by several laboratories researching cancer biology mechanisms, cancer biology mechanisms, neuromuscular junction function and Schwann cell proliferation [16]. NRG-1 has been previously demonstrated by our lab and others to be neuroprotective following ischemia in rat models [7-15]. We have shown that the exogenous delivery of NRG-1 reduces neuronal damage in the penumbral region of rats that underwent middle cerebral artery occlusion in an extended therapeutic window [10, 14, 15, 17-20]. While NRG-1 has shown significant promise in preventing brain damage and stimulating post-injury repair following stroke [21, 22], the regulatory mechanisms behind its neuroprotective effects are unclear. The investigation of ischemia-induced gene expression profiles following NRG-1 administration will provide valuable insight into the transcriptional regulation of these processes.

Previous studies have shown that ischemic stroke causes a release of pro-inflammatory cytokines that produce changes in gene expression, primarily in inflammation and cell death [4-6]. The initial area of injury, known as the infarct core, develops within minutes of stroke onset and develops up to three hours after stroke. The infarct core is characterized by low cerebral blood flow, oxidative stress, and excitotoxicity, which is accompanied by an increased production of inflammatory molecules. The resulting inflammation begins to affect a larger area of brain tissue, known as the ischemic penumbra, where neurons can survive up to 24-hours after stroke onset, prolonging the therapeutic window for ischemic stroke treatment [23].

In this study, we performed a computational transcriptomic analysis to identify gene expression profiles and their associated transcriptional mechanisms affected by NRG-1 treatment twelve hours following pMCAO using a transcriptome analysis software. To investigate mechanisms in the cortex following ischemia, we identified genes that were significantly regulated following NRG-1 treatment at twelve hours following ischemic stroke and the transcriptional pathways that regulated the expression of these genes. The PI3K-AKT pathway and its CREB1 and FOXO1 transcription factors have been recently reported to have a key role in ischemic stroke progression. We hypothesized that transcription factors related to ischemia-induced processes, such as the CREB1 and FOXO1 transcription factors, would be regulated with NRG-1 treatment. Understanding the mechanisms regulated by the NRG-1 phosphorylation cascade will help elucidate the neuroprotective effects of NRG-1 at the molecular level. These findings could support the development of clinical studies using NRG-1 to treat patients with ischemic stroke.

## Materials and Methods

### Animals

All animals were treated humanely with regard for the alleviation of pain and suffering. All surgical protocols involving animals were performed by aseptic techniques and were approved by the Institutional Animal Care and Use Committee at Morehouse School of Medicine prior to the experiment. Male adult Sprague-Dawley *rattus norvegucis* (250-300g; Charles River Laboratory International, Inc., USA) were housed in standard cages in a temperature-controlled room at 22 ± 2°C on a twelve-hour reverse light-dark cycle, where food and water were provided *ad libitum*.

Rats were randomly allocated into three groups: Sham (control), MCAO + vehicle treatment (MCAO), and MCAO + NRG-1 treatment (MCAO+NRG-1). All rats were anesthetized with 5% isoflurane with a O_2_/N_2_O mixture at 30%/70%. Once anesthetized, a rectal probe monitored the core body temperature of the animals, and a homoeothermic blanket control unit (Harvard Apparatus, Hollister, MA) was used to maintain a stable body temperature of 37°C. Cerebral blood flow was monitored during surgery by a continuous laser Doppler flowmeter (Perimed, Ardmore, PA). The Doppler probe was placed 7 mm lateral and 2 mm posterior to the bregma in a thinned cranial skull window.

Rats in the treatment groups (MCAO and MCAO+NRG-1) underwent a left pMCAO. MCAO was induced by the intraluminal suture method previously described [18]. These rats were then administered 50 μl of NRG-1β (1% BSA in PBS) or 50 μl of vehicle (1% BSA in PBS) treatment (20 ug/kg; EGF-like domain, R&D Systems, Minneapolis, MN) via bolus injection into the ICA through the ECA immediately before MCAO, as previously described [18]. The MCAO procedure involves inserting a 4 cm length 4–0 surgical monofilament nylon suture coated with silicon (Doccol Corp., Sharon, MA) from the external carotid artery (ECA) into the internal carotid artery (ICA) and then into the Circle of Willis to permanently occlude the left middle cerebral artery (MCA). Animals in the treatment groups were sacrificed twelve hours after MCAO. Rats in the control group (Sham) underwent a similar procedure as those in the treatment groups except a 4-0 surgical monofilament was not inserted into the ICA. Rats in the control group were sacrificed three hours after the Sham procedure. A subset of the brains were used for 2,3,5-triphenyltetrazolium chloride (TTC) staining as previously described [18]. With TTC labeling, the infarcted region appears white, whereas the normal non-infarcted tissue appears red. All NRG-1 and vehicle treatment studies were performed in a blinded manner.

### Microarray Analysis

Treated animals were sacrificed twelve hours following MCAO (n = 4 for MCAO and n = 5 for MCAO+NRG-1) and control animals were sacrificed three hours after Sham surgery (n = 4). Brains were extracted and sectioned into 2 mm coronal sections, approximately +3.0 to −5.0 from bregma, where left cortical tissue was isolated. Left hemi-cortical tissue from Sham was used as the control group. Total RNA was extracted with TRIzol Reagent (Life Technologies Corp, Carlsbad, CA), then controlled and quantified by the Agilent 2100 Bioanalyzer (Agilent Technology, Santa, Clara, CA). Microarrays (Affymetrix Inc., Santa, Clara, CA) were completed according to manufacturing guidelines, with mRNA hybridized to an Affymetrix Rat Genome 2.0st Gene Chip (Affymetrix Inc.). Microarray chips were then used for gene expression analysis.

Statistical analysis of microarray data was completed using the Affymetrix Transcriptome Analysis Console (TAC) 4.0 Software Package. The original *.CEL files generated from the gene chip were imported to TAC for quality control, data normalization, and to identify differentially expressed genes. The transcriptome analysis approach was utilized due to its ability to simultaneously analyze a large amount of gene changes and identify possible regulating gene groups. For differential expression analyses, a cutoff of 1.5-fold change and p-value of < 0.05 were used. Lists of differentially expressed genes between different conditions were saved as a .TXT file for further use in other software.

From the list of genes generated by the TAC software, transcription factors were identified with the JASPAR open-access database [24]. Of the identified transcription factors, the Fiserlab TF2DNA database [25] was used to find downstream regulated molecules in order to determine if the genes regulated by those factors experience expression changes observed in the microarray.

### Multiplex Transcription Factor Array

Brain tissues from the ipsilateral cortex, which are lateral to sections obtained for RNA extractions for the microarray analysis, were used for transcription factor assay studies (n = 3 for each condition). Nuclear proteins were isolated using the USB Nuclear Extraction Kit (Affymetrix, Inc.) and the Procarta Transcription Factor Assay (Affymetrix, Inc.) was used to measure transcription factor DNA binding activity. All protocols were performed according to the manufacturer’s guidelines.

The Procarta detection probes and nuclear extracts from the samples were incubated for thirty minutes at 15°C to create a protein-DNA complex, and Procarta controls (n = 3) and blanks (n = 3) were added. All samples were washed with a cold binding buffer on a 96-well separation plate to remove unbound protein and DNA, and were then incubated on ice once for five minutes, centrifuged for five minutes at 4°C, and the flow through was discarded. Centrifugation and flow discard was repeated five times. Samples were heated to 95°C for five minutes to denature the Protein-DNA complex and were then incubated with capture beads for 30 minutes at 50°C on a shaker for 400 rpm. Streptavidin-PE was added to the capture beads for thirty minutes at room temperature on a shaker for 400 rpm before the beads were washed and resuspended in reading buffer on a shaker at 400 rpm. Results were read on the Luminex Bio-plex system 200 (Bio-Rad, Hercules, California). Activity levels of transcription factors identified with JASPAR were examined and reported.

### Protein-Protein Interaction Network Analysis

The Search Tool for the Retrieval of Interacting Genes/Proteins (STRING) database (16) was used to identify protein-protein interactions between the genes regulated in the microarray, as well as determine the relationship between the transcriptional factors found through JASPAR. STRING allows users to obtain a visualized network of protein-protein interactions from a list of proteins uploaded to the website, where each node corresponds to a gene and genes are connected by lines whose thickness reflects the strength of data evidence. STRING interaction confidence scores are generated by collecting multiple types of data (such as prior co-expression, experimental data, association in curated databases, and more) to determine the probability of an interaction.

A list of all genes that were upregulated between MCAO and NRG-1 treatments in the microarray data were inputted into STRING and the connections between genes within our data set were visualized. Any orphan gene that was not associated with the network was removed from view and further analysis. Similarly, a list of transcription factors identified by JASPAR and examined using the transcription factor array were uploaded into STRING, and the first five interactors were mapped to view a network expansion and identify the relationship between the factors.

## Results

### Microarray Analysis

We previously demonstrated neuroprotection by NRG-1 following ischemia in rat models [7-15]. In these studies, rats were treated with NRG-1 immediately before MCAO and brain tissues were collected 12 hours following ischemia. TTC staining detected large infarcts in brain tissues 12 hours following MCAO (Fig 1A) which was reduced by NRG-1 treatment (Fig 1B) as previously shown when measure 24 hours following MCAO. Here, we examined gene expression profiles in the ischemic brain cortex 12 hours following vehicle and NRG-1 treatment. Of the total 30,429 gene probes on the Affymetrix Rat 2.0st chip, 21,526 genes were annotated for the rat genome. The microarray plots in figure 2 show the relationship between the signal intensity for the expressed genes and difference between two conditions for those genes. Following MCAO, there was a significant increase in the number of genes upregulated and downregulated compared to Sham. The plots in figure 2A compare the expression intensities of the 1350 genes differentially expressed between the Sham control and MCAO. The plots in figure 2B compare the intensities of the 1979 genes differentially expressed for the Sham control and MCAO+NRG-1 conditions, featuring a larger increase in the number of genes being upregulated and downregulated (Fig 2A). When comparing gene expression from the MCAO condition to rats treated with NRG-1 before MCAO (Fig 2C), there were 556 genes which were regulated by NRG-1. Of the 556 genes, 253 genes were upregulated, as shown in the upper red dots in figure 2C and the right red dots in the volcano plot in figure 2D. The 253 genes that were upregulated between the MCAO and NRG-1 groups with a fold change (fc) > 1.5 and p-value < 0.05 were selected for further analysis.

**Figure 1.**
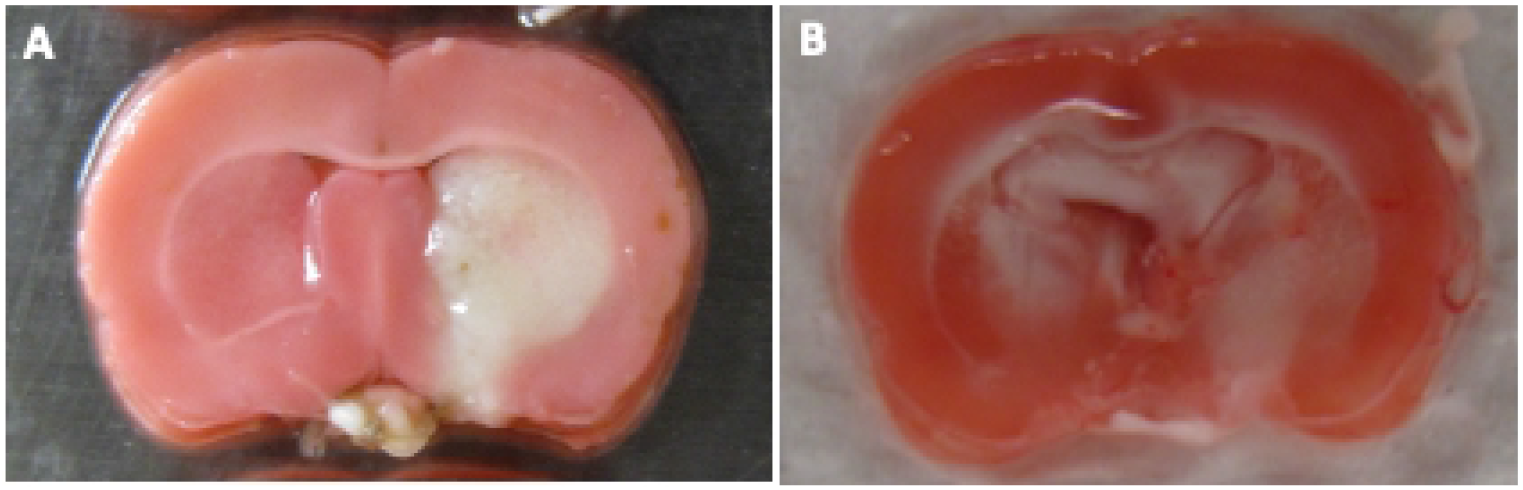
Neuronal Injury 12 Hours Following MCAO. NRG-1 reduced infarct volume following MCAO. Mice were treated with NRG-1 or vehicle immediately after MCAO. Brains were removed 12 hours after MCAO, sliced into coronal sections (2 mm thick), and treated with 2% TTC. Compared to vehicle treated animals (A), NRG-1 reduced neuronal injury (B).

**Figure 2.**
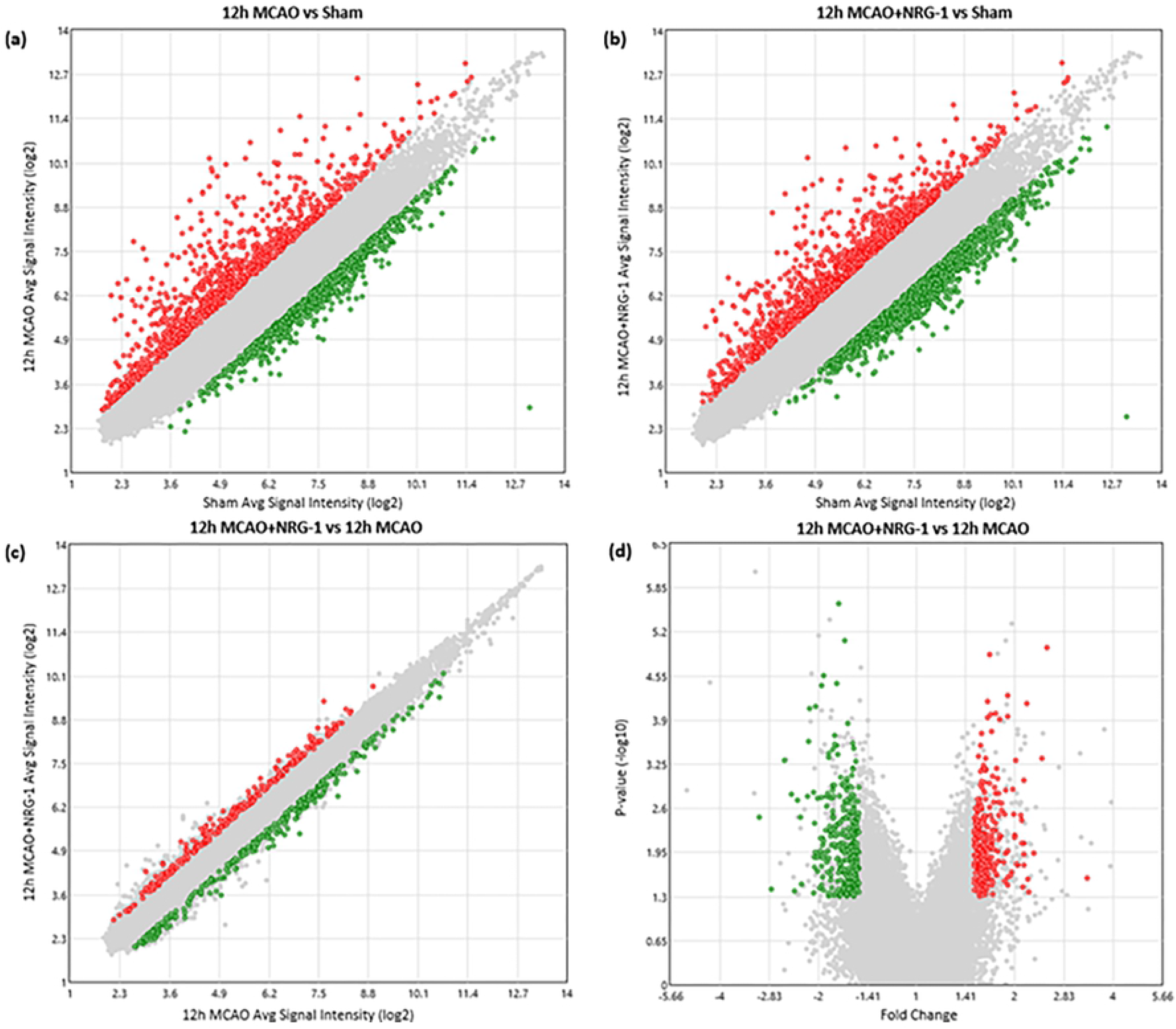
Differential Gene Expression Analysis. (a) 1350 genes were differentially expressed between the Sham and 12 h MCAO conditions, showing differences in gene expression due to injury. Red dots (upper left side) indicate genes that were upregulated between these conditions and green dots (lower right side) indicate genes that were downregulated between conditions. (b) 1979 genes were differentially expressed between the Sham and 12 h MCAO+NRG-1 conditions. Red dots (upper left side) indicate genes that were upregulated between these conditions and green dots (lower right side) indicate genes that were downregulated between conditions. (c) 556 genes were differentially expressed between the 12h MCAO and 12 h MCAO+NRG-1 conditions, depicting the changes in gene expression caused by NRG-1 delivery after injury. Red dots (upper left side) indicate genes that were upregulated between these conditions and green dots (lower right side) indicate genes that were downregulated between conditions. (d) Volcano plot shows the p-value vs fold change of genes in Fig 1C. Green dots (upper left side) indicate genes that were significantly downregulated between these conditions and red dots (upper right side) indicate genes that were upregulated between conditions.

In Table 1, genes that underwent separate expression patterns were categorized and clustered based on statistical expression changes. There were 26 genes that increased expression with MCAO (fc > 1.5) and further increased with NRG-1 (fc > 1.5), which are associated with metabolic processes, cellular growth, biological regulation, stimuli response, and localization. These genes are in pathways such as CCKR signaling, cadherin signaling, and cytokine mediated inflammation. There were 188 genes whose expression remained unaltered following MCAO (−1.5 < fc < 1.5) but then increased following NRG-1 (fc > 1.5) treatment were associated with metabolic processes, cellular growth, biological regulation, and immune system function. These genes are in pathways such as oxidative stress response, p53 signaling, DNA replication, PI3K pathway, EGF signaling, and more. Of these genes, three are transcription factors: CREB1, HIC2, and ZBTB26. There were 39 genes that decreased expression with MCAO (fc < -1.5) and were rescued with NRG-1 (fc > 1.5), which are associated with metabolic process, cellular development, cell maintenance, and biological regulation. These genes are in pathways such as CCKR signaling, cadherin signaling, and Insulin/IGF pathway, PI3K pathway, and Wnt signaling. Of these genes, two are transcription factors: ZIC2 and FOXO1.

**Table 1.**
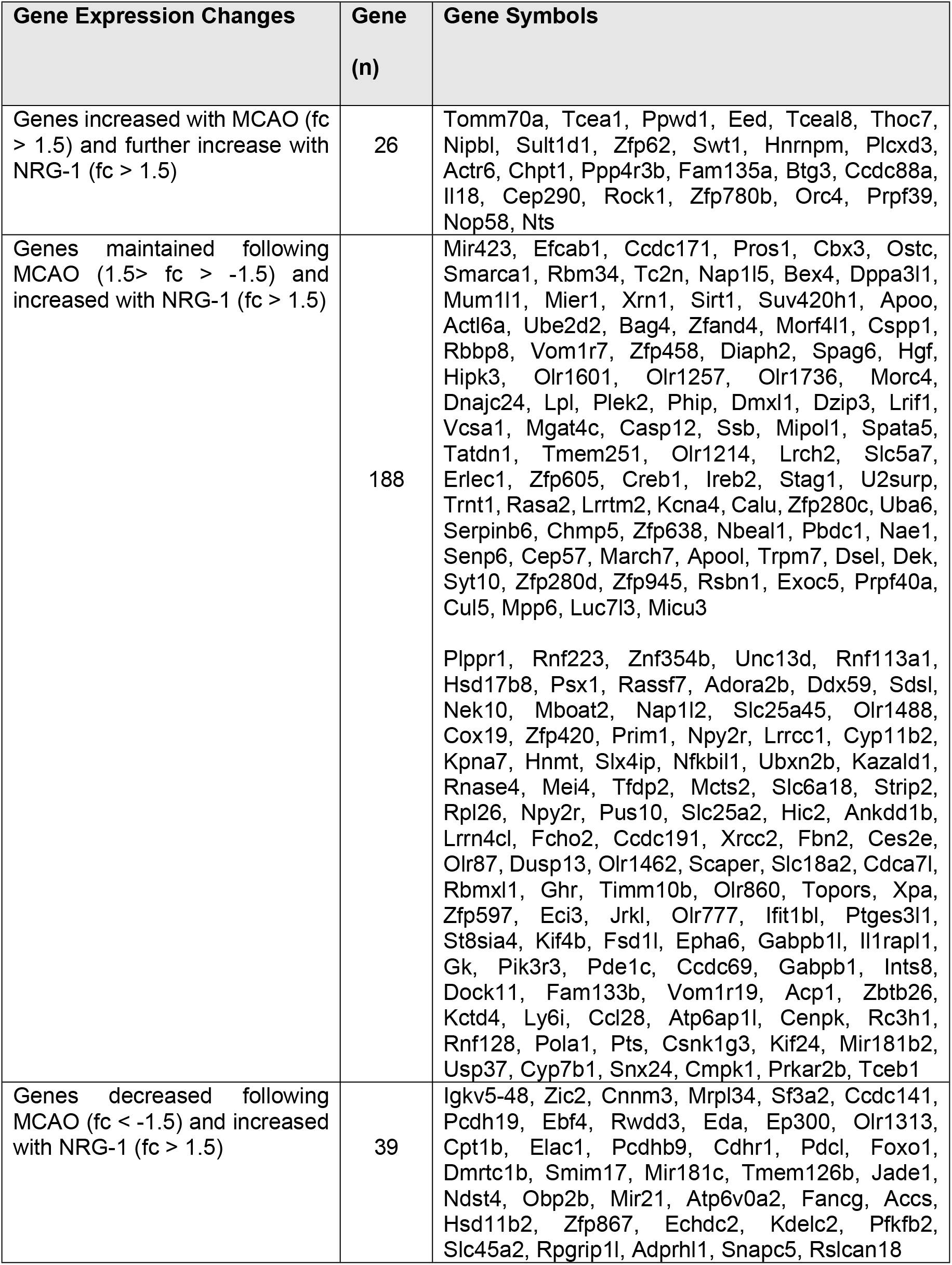
Genes with change from Sham due to MCAO and subsequent significant upregulation due to NRG-1 after 12 hours.

The 253 genes listed in Table 1 were uploaded into the STRING database to understand the relationships between them, their roles related to ischemic stroke, and identify key regulatory factors. The thickness of the connecting lines reflects the confidence interval of the interaction, with darker lines signifying high confidence that the proteins these genes encode interact with one another. The number of connecting nodes indicate the number of interactors the gene has, where a gene with a high number of interactors may signify a potential regulator.

Figure 3 shows the network for genes upregulated with fc > 1.5 between the 12h MCAO and 12h MCAO+NRG-1 conditions and orphan genes unconnected to the network were removed from view. Of the 5 transcription factors originally identified in the microarray analysis, only CREB1, FOXO1, and ZBTB26 were connected to the network. There were three main clusters of genes that were identified. Hub 1 contains the CREB1 and FOXO1 transcription factors (Fig 3). The ZBTB26 transcription factor was not featured in a hub, and no transcription factors were present in either hubs 2 or 3. Due to containing multiple transcription factors, Hub 1 was selected for future transcription factor analysis.

**Figure 3.**
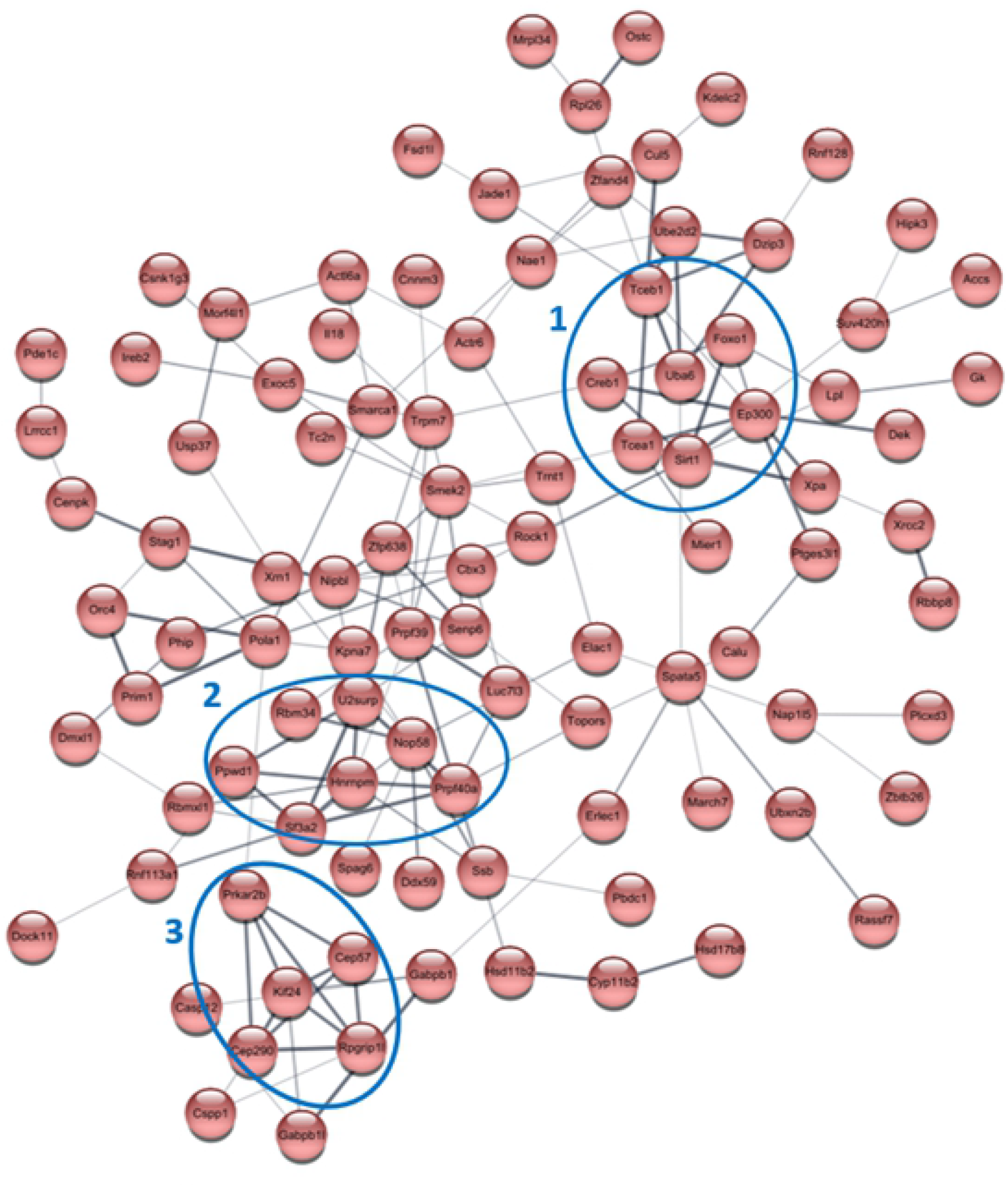
Network interaction of genes (n = 99) upregulated with fc > 1.5 between the 12h MCAO and 12h NRG conditions. Protein-protein interactions visualized using STRING are used to identify potential “hub” genes that signify regulatory factors and related pathways. The thickness of the connecting line represents the confidence interval of the interaction. Orphan genes unconnected to the network were removed from view. Of the 5 transcription factors identified in the microarray analysis, only CREB1, FOXO1, and ZBTB26 were connected to the network. Circled are 3 potential “hubs” for analysis, indicated as hubs 1,2, and 3. Notably, the CREB1 and FOXO1 transcription factors are closely connected and are featured in Node 1, highlighting a potentially significant “hub” and related pathway involving these genes. The ZBTB26 transcription factor was not featured in a hub, and no transcription factors were present in hubs 2 or 3, therefore Node 1 was selected for further analysis.

When examining the downstream targets of the identified CREB1 and FOXO1 transcription factors using the FiserLab TF2DNA database, several targets were seen to be regulated by MCAO and NRG-1 treatment in our dataset. Downstream targets of FOXO1 that were present in the list of genes from our microarray dataset include PPWD1, DCAF4, TATDN1, SLC25A12, and NDST4. The microarray expression patterns of CREB1 and FOXO1 and their downstream targets are shown in Figure 4. FOXO1 and NDST4 exhibited downregulation after MCAO and rescue by NRG-1. CREB1, PWWD1 and TATDN1 were characterized by slight increases following MCAO and further increases following NRG-1 treatment. DCAF4 and SLC24A2 shown downregulation after MCAO and were further decrease by NRG-1.

**Figure 4.**
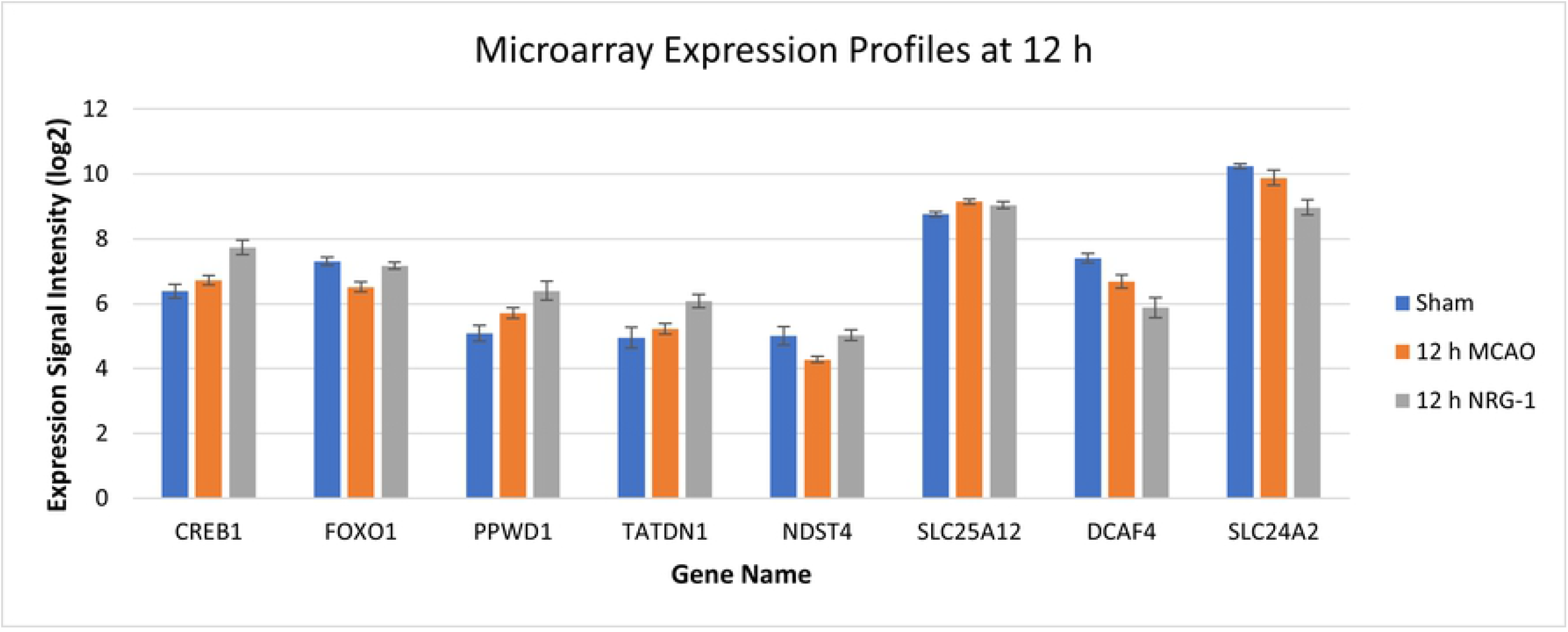
Gene expression profiles among statistically significant genes (ANOVA, p < 0.05) in Sham (n=4), MCAO (n=4), and MCAO+NRG-1 (n=5) treated animals at 12 h following injury. CREB1 and FOXO1 transcription factors were identified to be significantly upregulated with NRG-1 compared to MCAO. Downstream target of CREB1 that was regulated by NRG-1 compared to MCAO is SLC24A2, which followed a descending trend. Downstream targets of FOXO1 that were regulated by NRG-1 compared to MCAO are PPWD1, TATDN1, NDST4, SLC25A12, and DCAF4, where the first three genes are present in Table 1.

### CREB1 and FOXO1 Transcription Factor Network Analysis

The FOXO1 and CREB1 transcription factors identified through microarray analysis and examined in the transcription factor array displayed high confidence interactions with one another. Interested in the relationship of these two factors, the STRING database was used to map the first five notable interactors of the CREB1 and FOXO1. The immediate interactors identified by STRING are CREBBP, SIRT1, CRTC2, AKT1, and ATF1, as shown in an expansion of the network (Fig 5). Of the five interactors, SIRT1 was shown to be upregulated due to NRG-1 in our dataset. Additionally, several of these genes are featured in the PI3K-AKT pathway, highlighting a common pathway between CREB1 and FOXO1 that is not only an important mechanism in ischemic stroke, but suggests that NRG-1 assists in recovery through the PI3K-AKT pathway.

**Figure 5.**
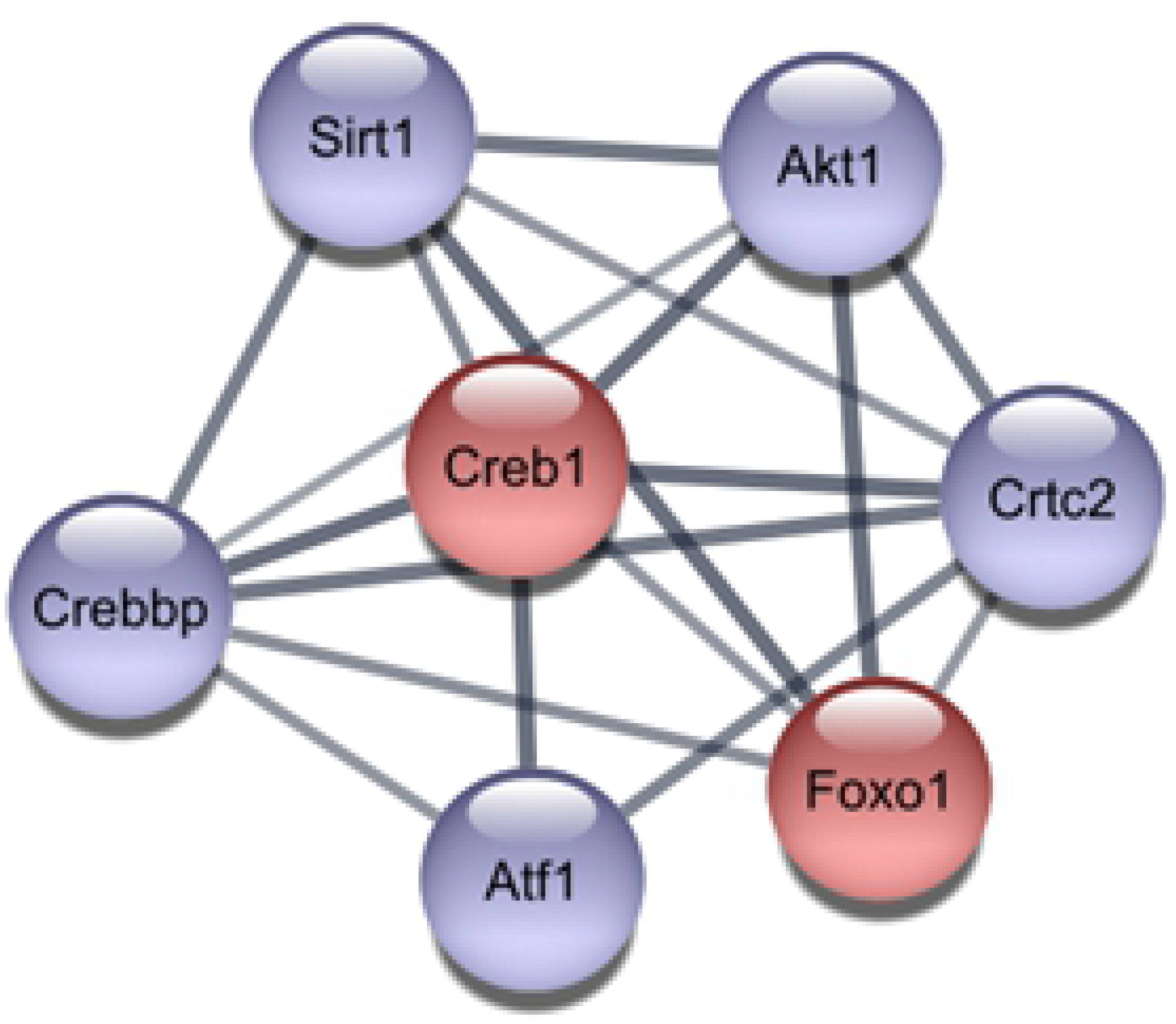
Network expansion of the CREB1 and FOXO1 transcription factors. Protein-protein interactions of the upregulated transcription factor of interest (CREB1 and FOXO1, shown in red) were visualized using STRING and the network was expanded to include the first 5 interactors (shown in purple). Network expansion was performed to identify potential pathways involved with the CREB1 and FOXO1 transcription factors.

### Multiplex Transcription Factor Array

To examine whether the predicted transcriptional regulators were involved with ischemia and NRG-1 neuroprotection, we conducted multiplex transcription factor assays on nuclear extracts from the brains of Sham, MCAO, and MCAO+NRG1 animals. FOXO1 and CREB were present on the transcription factor array plate and both transcription factors featured significant activity changes with NRG-1 delivery 12 hours after stroke (Fig 6). FOXO1 exhibited an increase of activity following MCAO and further increased with NRG-1 treatment. CREB1 displayed decreased activity after MCAO but was rescued with NRG-1 treatment. The increased activity of CREB1 and FOXO1 by NRG-1 and the increased microarray expression of these factors following NRG-1 treatment suggest that these factors are involved in the neuroprotective effects of NRG-1.

**Figure 6.**
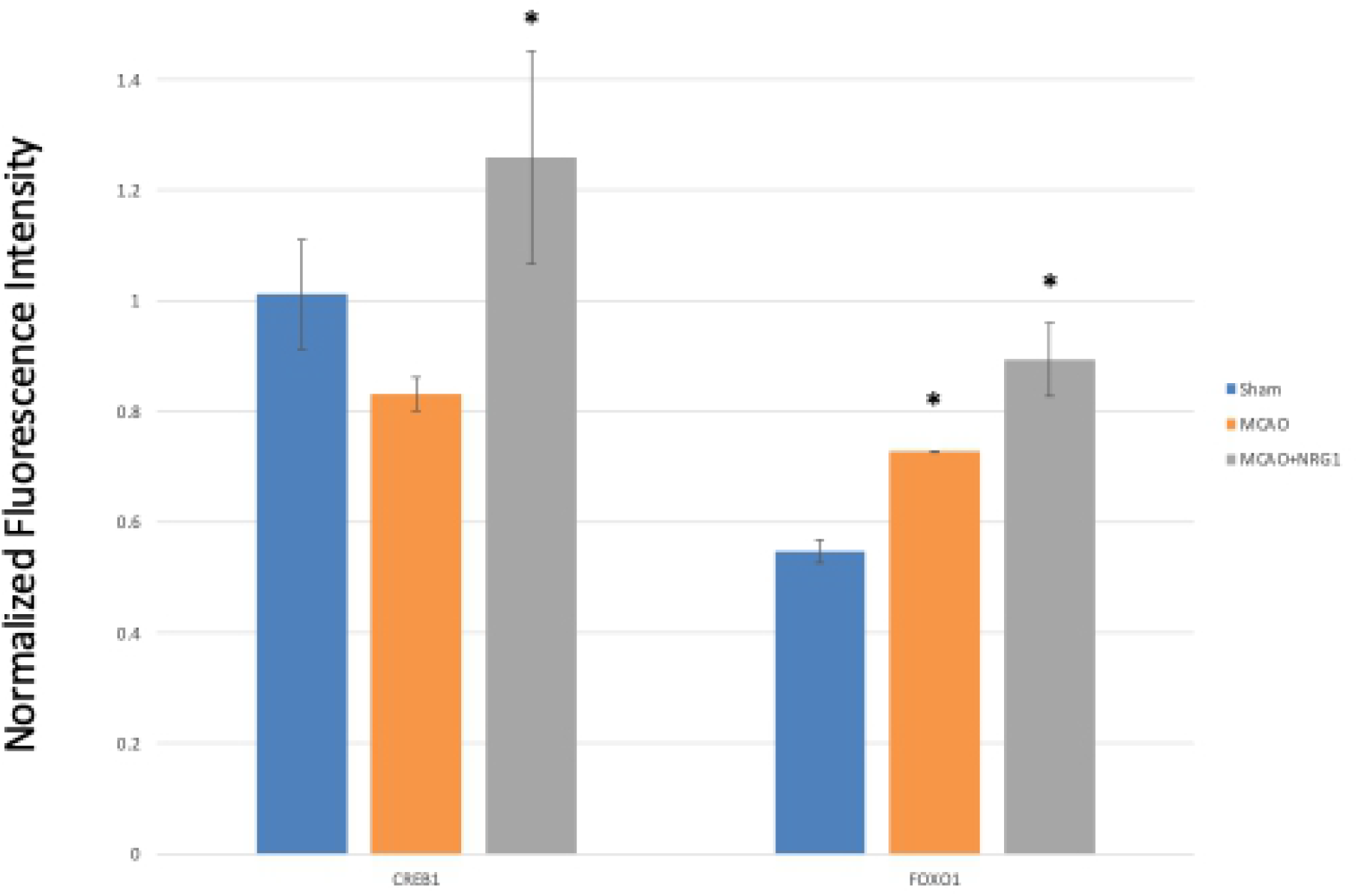
The activity level of the CREB1 and FOXO1 transcription factors. With NRG-1 treatment 12 h after injury, transcription factor activity of CREB1 and FOXO1 increase compared to MCAO. CREB1 was downregulated due to ischemic stroke, and NRG-1 treatment rescued CREB1 activity. FOXO1 activity was slightly upregulated after ischemic stroke, and NRG-1 treatment further promoted FOXO1 activity.

## Discussion

NRG-1 has demonstrated neuroprotective and anti-inflammatory properties in rats and mice following ischemic stroke [7-15], however, the exact molecular mechanisms involved in NRG-1 neuroprotection remains unclear. We previously showed that the neuroprotective and anti-inflammatory effects of NRG-1 in ischemia involve the activation of PI-3 kinase (PI3K)-Akt pathways and inhibition of canonical NF-κB mediated signaling [26]. The current study expands on these findings based on microarray and transcription factor array data taken from infarct tissue at 12 hours post-stroke to further elucidate the signaling and transcriptional mechanisms involved in the neuroprotective effects of NRG-1. Of the total 30,463 genes analyzed, 565 genes were significantly differentially expressed between the MCAO and MCAO+NRG-1 groups. Within that group, we identified CREB1 and FOXO1 transcription factors as potential key regulatory of NRG-1 mediated effects following ischemia. The CREB1 and FOXO1 transcription factors have been shown to play roles in cell survival following ischemic stroke that are heavily interconnected and dependent upon one another. CREB1, a cAMP responsive element binding protein activated by the ep300 protein, is a transcription factor that drives several cellular responses, such as cell proliferation, survival, and differentiation [27]. CREB1 has been identified in having specific immune functions, including inhibiting canonical NF-kB pro-inflammatory activation, inducing macrophage survival, and promoting the proliferation, survival, and regulation of T and B lymphocytes [27]. Neuroprotection by equilibrative nucleoside transporter one (ENT1) was shown to be mediated by increased activity of CREB associated with decreased neuronal apoptosis [28, 29]. Other neuroprotective compounds including adiponectin and zeranol have been shown to reduce ischemia-induced neuronal injury by activating CREB signaling pathways [30]. The upregulation of ep300 suggests that CREB is being activated, leading to neuronal survival.

Forkhead box class O (FOXO) proteins are transcription factors that have been shown to play roles in a number of neurological disorders including Alzheimer’s disease, Parkinson’s disease and amyotrophic lateral sclerosis [31, 32]. FOXO proteins interact with a number of signaling pathways including Akt and SIRT1. FOXO1 and FOXO3 have been reported to regulate apoptosis, oxidative stress and autophagy in neuronal cells downstream of Akt and SIRT1. Estradiol reduces neuronal death following ischemia and prevents ischemia-induced decrease of FOXO1 and Akt phosphorylation [33].

Additionally, the expression level of SIRT1, a nicotinamide adenosine dinucleotide-dependent histone deacetylase downstream of the PI3K-AKT pathway, was upregulated in this dataset. SIRT1 is a transcriptional co-factor that plays a role in longevity, apoptosis, and stress resistance by deacetylating proteins, such as p53, NF-κB, and FOXO transcription factors [34]. SIRT1 was shown to promote FOXO1-induced autophagy and cell cycle arrest while decreasing apoptosis following ischemia/hypoxia in multiple cell types [35, 36]. Through deacetylation, SIRT1 has also been shown to regulate the transcriptional activity of NF-κB and protect cells from p53-induced apoptosis [37, 38]. Sirt1-overexpressing mice were shown to protect against cerebral global ischemia [39].

The model in figure 7 illustrated how we propose that NRG-1 prevents neuronal death and neuroinflammation following ischemic stroke. NRG-1 binds to erbB4 receptors which signal by activating the PI3K-Akt pathway activating CREB and increasing neuronal survival. Upregulation of FOXO1 by NRG-1 can prevent oxidate stress and apoptosis and also protect neurons. Upregulation of SIRT1 by NRG-1 can result in the inhibition of NF-κB leading to anti-inflammatory effects following ischemia.

**Figure 7.**
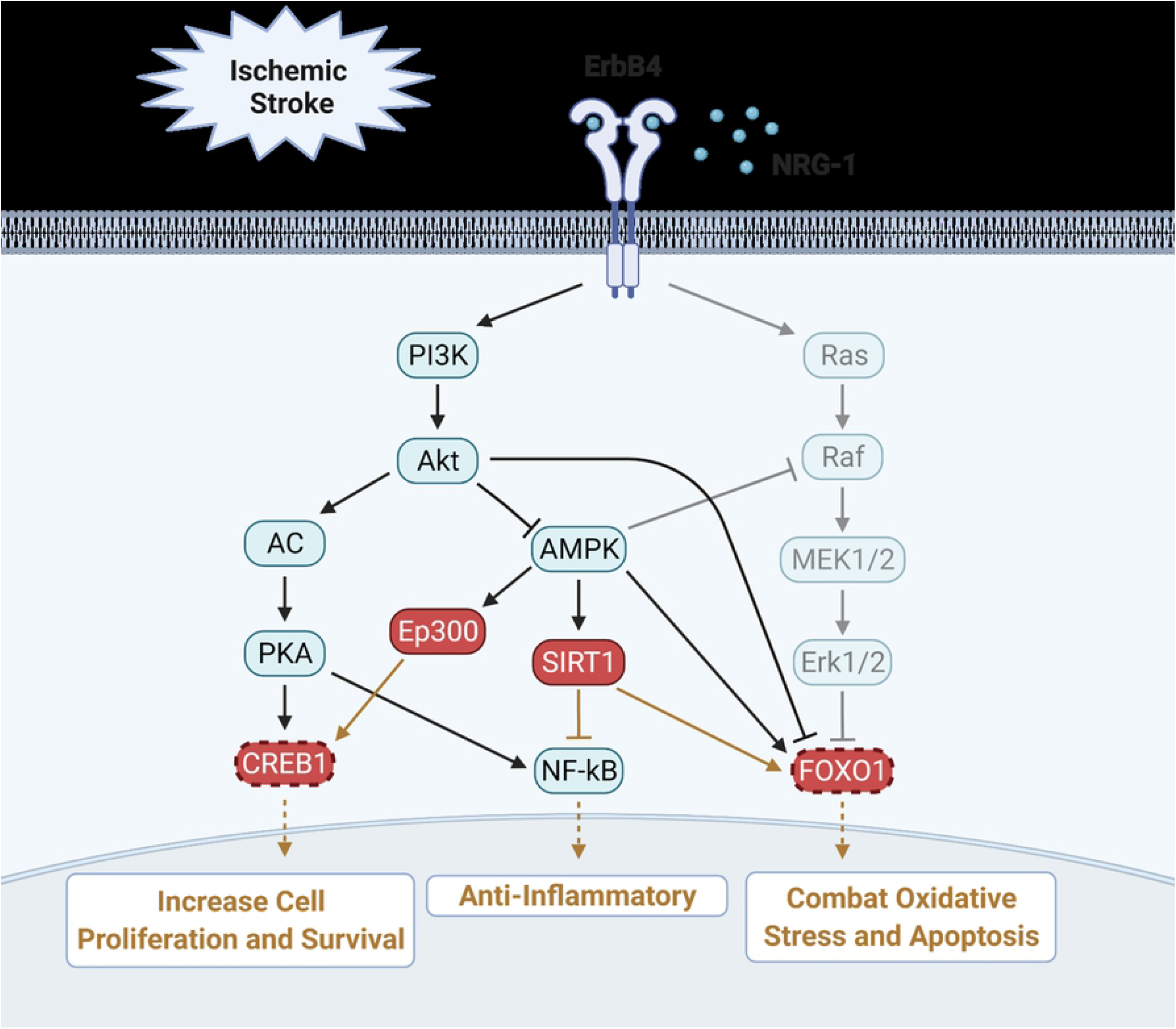
Suggested pathway analysis of NRG-1 neuroprotection and involvement in the PI3K-AKT pathway. Genes upregulated with NRG-1 delivery compared to MCAO are shown in red, transcription factors are shown in red with dotted outlines, black arrows are inferred from literature, and proposed effects due to gene upregulation are shown in brown coloration. Upregulation of ep300 and Sirt1 lead to the upregulation of the CREB1 and FOXO1 transcription factors. Regulation of both two transcription factors suggest stimulating effects of NRG-1 delivery on the PI3K-AKT pathway. Downstream effects of this pathway have been associated with pro-survival and anti-inflammatory response, highlighting potential outcomes of NRG-1 delivery in ischemic stroke. Figure created using BioRender.com.

## Conclusion

We have previously shown that NRG-1 is neuroprotective and anti-inflammatory following ischemic stroke. CREB and FOXO1 were identified as a candidate transcription factor pathways associated with the regulation of gene expression by stroke and NRG-1 treatment. NRG-1 isoforms were shown to improve cardiac function in clinical trials of human patients [40]. Therefore, understanding the cellular and molecular mechanisms of neuroprotection by NRG-1 could lead to the development of a novel stroke treatment and to the identification of novel targets for therapeutic intervention.

## Author Contributions

**Conceptualization**: Kimberly Bennett, Catherine Augello, Victor Rodgers, Byron Ford

**Data curation:** Monique Surles-Zeigler

**Formal analysis:** Kimberly Bennett, Catherine Augello, Etchi Ako

**Funding acquisition:** Kimberly Bennett, Etchi Ako, Victor Rodgers, Byron Ford

**Investigation:** Kimberly Bennett, Catherine Augello, Etchi Ako

**Methodology:** Monique Surles-Zeigler, Byron Ford

**Project administration:** Victor Rodgers, Byron Ford

**Resources:** Byron Ford

**Software:** Kimberly Bennett, Catherine Augello

**Supervision:** Victor Rodgers, Byron Ford

**Validation:** Monique Surles-Zeigler, Byron Ford

**Visualization**: Kimberly Bennett, Catherine Augello

**Writing – original draft:** Kimberly Bennett

**Writing – review & editing:** Kimberly Bennett, Catherine Augello, Victor Rodgers, Byron Ford

## Data Availability

The data will be submitted to Gene Expression Omnibus (https://www.ncbi.nlm.nih.gov/geo/) before publication.

## References

1. Fisher, M., et al., Update of the Stroke Therapy Academic Industry Roundtable Preclinical Recommendations. Stroke, 2009. 40: p. 2244–2250.

2. Bosetti, F., et al., Translational Stroke Research: Vision and Opportunities. Stroke, 2017. 48(9): p. 2632–2637.

3. Benjamin, Heart Disease and Stroke Statistics-2017 Update: A Report From the American Heart Association (vol 135, pg e146, 2017). Circulation, 2017. 136(10): p. E196–E196.

4. Jayaraj, R.L., et al., Neuroinflammation: friend and foe for ischemic stroke. J Neuroinflammation, 2019. 16(1): p. 142.

5. Dirnagl, U., C. Iadecola, and M.A. Moskowitz, Pathobiology of ischaemic stroke: an integrated view. Trends Neurosci, 1999. 22(9): p. 391–7.

6. Barone, F.C. and G.Z. Feuerstein, Inflammatory mediators and stroke: new opportunities for novel therapeutics. J Cereb Blood Flow Metab, 1999. 19(8): p. 819–34.

7. Surles-Zeigler, M.C., et al., Transcriptomic analysis of neuregulin-1 regulated genes following ischemic stroke by computational identification of promoter binding sites: A role for the ETS-1 transcription factor. Plos One, 2018. 13(6).

8. Xu, Z., et al., Neuroprotection by neuregulin-1 following focal stroke is associated with the attenuation of ischemia-induced pro-inflammatory and stress gene expression. Neurobiol Dis, 2005. 19(3): p. 461–70.

9. Ford, G., et al., Expression Analysis Systematic Explorer (EASE) analysis reveals differential gene expression in permanent and transient focal stroke rat models. Brain Res, 2006. 1071(1): p. 226–36.

10. Xu, Z., et al., Neuregulin-1 is neuroprotective and attenuates inflammatory responses induced by ischemic stroke. Biochem Biophys Res Commun, 2004. 322(2): p. 440–6.

11. Rodriguez-Mercado, R., et al., Acute Neuronal Injury and Blood Genomic Profiles in a Nonhuman Primate Model for Ischemic Stroke. Comparative Medicine, 2012. 62(5): p. 427–438.

12. Simmons, L.J., et al., Regulation of inflammatory responses by neuregulin-1 in brain ischemia and microglial cells in vitro involves the NF-kappa B pathway. Journal of Neuroinflammation, 2016. 13.

13. Wang, S.L., et al., Spatio-temporal assessment of the neuroprotective effects of neuregulin-1 on ischemic stroke lesions using MRI. Journal of the Neurological Sciences, 2015. 357(1-2): p. 28–34.

14. Xu, Z., et al., Extended therapeutic window and functional recovery after intraarterial administration of neuregulin-1 after focal ischemic stroke. J Cereb Blood Flow Metab, 2006. 26(4): p. 527–35.

15. Noll, J.M., et al., Neuroprotection by Exogenous and Endogenous Neuregulin-1 in Mouse Models of Focal Ischemic Stroke. J Mol Neurosci, 2019. 69(2): p. 333–342.

16. Cespedes, J.C., et al., Neuregulin in Health and Disease. Int J Brain Disord Treat, 2018. 4(1).

17. Shyu, W.C., et al., Neuregulin-1 reduces ischemia-induced brain damage in rats. Neurobiology of Aging, 2004. 25(7): p. 935–944.

18. Li, Y., et al., Neuroprotection by neuregulin-1 in a rat model of permanent focal cerebral ischemia. Brain Res, 2007. 1184: p. 277–83.

19. Li, Q., et al., Neuregulin attenuated cerebral ischemia-Creperfusion injury via inhibiting apoptosis and upregulating aquaporin-4. Neuroscience Letters, 2008. 443(3): p. 155–159.

20. Guan, Y.F., et al., Neuregulin 1 protects against ischemic brain injury via ErbB4 receptors by increasing GABAergic transmission. Neuroscience, 2015. 307: p. 151–9.

21. Iaci, J.F., et al., Glial growth factor 2 promotes functional recovery with treatment initiated up to 7 days after permanent focal ischemic stroke. Neuropharmacology, 2010. 59(7-8): p. 640–9.

22. Iaci, J.F., et al., An optimized dosing regimen of cimaglermin (neuregulin 1β3, glial growth factor 2) enhances molecular markers of neuroplasticity and functional recovery after permanent ischemic stroke in rats. Journal of Neuroscience Research, 2016. 94(3): p. 253–265.

23. Lipton, P., Ischemic cell death in brain neurons. Physiol Rev, 1999. 79(4): p. 1431–568.

24. Fornes, O., et al., JASPAR 2020: update of the open-access database of transcription factor binding profiles. Nucleic Acids Res, 2020. 48(D1): p. D87–D92.

25. Pujato, M., et al., Prediction of DNA binding motifs from 3D models of transcription factors; identifying TLX3 regulated genes. Nucleic Acids Res, 2014. 42(22): p. 13500–12.

26. Simmons, L.J., et al., Regulation of inflammatory responses by neuregulin-1 in brain ischemia and microglial cells in vitro involves the NF-kappa B pathway. J Neuroinflammation, 2016. 13(1): p. 237.

27. Wen, A.Y., K.M. Sakamoto, and L.S. Miller, The role of the transcription factor CREB in immune function. J Immunol, 2010. 185(11): p. 6413–9.

28. Zhang, D., et al., ENT1 inhibition attenuates apoptosis by activation of cAMP/pCREB/Bcl2 pathway after MCAO in rats. Exp Neurol, 2020. 331: p. 113362.

29. Mohamed, S.K., et al., The protective effect of zeranol in cerebral ischemia reperfusion via p-CREB overexpression. Life Sci, 2019. 217: p. 212–221.

30. Bai, H., et al., Adiponectin confers neuroprotection against cerebral ischemia-reperfusion injury through activating the cAMP/PKA-CREB-BDNF signaling. Brain Res Bull, 2018. 143: p. 145–154.

31. Maiese, K., FoxO proteins in the nervous system. Anal Cell Pathol (Amst), 2015. 2015: p. 569392.

32. Maiese, K., Targeting the core of neurodegeneration: FoxO, mTOR, and SIRT1. Neural Regen Res, 2021. 16(3): p. 448–455.

33. Jover-Mengual, T., et al., Acute estradiol protects CA1 neurons from ischemia-induced apoptotic cell death via the PI3K/Akt pathway. Brain Res, 2010. 1321: p. 1–12.

34. Chae, H.D. and H.E. Broxmeyer, SIRT1 deficiency downregulates PTEN/JNK/FOXO1 pathway to block reactive oxygen species-induced apoptosis in mouse embryonic stem cells. Stem Cells Dev, 2011. 20(7): p. 1277–85.

35. Meng, X., et al., Sirt1: Role Under the Condition of Ischemia/Hypoxia. Cell Mol Neurobiol, 2017. 37(1): p. 17–28.

36. Chen, C.J., et al., Resveratrol protects cardiomyocytes from hypoxia-induced apoptosis through the SIRT1-FoxO1 pathway. Biochem Biophys Res Commun, 2009. 378(3): p. 389–93.

37. Yeung, F., et al., Modulation of NF-kappaB-dependent transcription and cell survival by the SIRT1 deacetylase. EMBO J, 2004. 23(12): p. 2369–80.

38. Brunet, A., et al., Stress-dependent regulation of FOXO transcription factors by the SIRT1 deacetylase. Science, 2004. 303(5666): p. 2011–5.

39. Hattori, Y., et al., SIRT1 attenuates severe ischemic damage by preserving cerebral blood flow. Neuroreport, 2015. 26(3): p. 113–7.

40. Meyer, D. and C. Birchmeier, Multiple essential functions of neuregulin in development. Nature, 1995. 378(6555): p. 386–390.

